# CTDSP1 inhibitor rabeprazole regulates DNA-PKcs dependent topoisomerase I degradation and irinotecan drug resistance in colorectal cancer

**DOI:** 10.1101/2020.01.07.897355

**Authors:** Hiroya Matsuoka, Koji Ando, Emma J Swayze, Elizabeth C Unan, Joseph Mathew, Quingjiang Hu, Yasuo Tsuda, Yuichiro Nakashima, Hiroshi Saeki, Eiji Oki, Ajit K Bharti, Masaki Mori

**Affiliations:** Department of Surgery and Science, Graduate School of Medical Sciences, Kyushu University, Fukuoka City, Fukuoka, Japan; Department of Medicine, Division of Hematology and Medical Oncology, Boston University School of Medicine, Boston, Massachusetts, United States of America; Department of General Surgical Science, Graduate School of Medicine, Gunma University, Maebashi, Gunma, Japan

## Abstract

Irinotecan specifically targets topoisomerase I (topoI), and is used to treat various solid tumors, but only 13-32% of patients respond to the therapy. Now, it is understood that the rapid rate of topoI degradation in response to irinotecan causes irinotecan resistance. We have published that the deregulated DNA-PKcs kinase cascade ensures rapid degradation of topoI and is at the core of the drug resistance mechanism of topoI inhibitors, including irinotecan. We also identified CTD small phosphatase 1 (CTDSP1) (a nuclear phosphatase) as a primary upstream regulator of DNA-PKcs in response to topoI inhibitors. Previous reports showed that rabeprazole, a proton pump inhibitor (PPI) inhibits CTDSP1 activity. The purpose of this study was to confirm the effects of rabeprazole on CTDSP1 activity and its impact on colon cancer. Using HCT116 and HT29, with high and low CTDSP1 expression respectively and a retrospective analysis of patients receiving irinotecan with or without rabeprazole have indicated the effect of CTDSP1 in irinotecan response. These results indicate that CTDSP1 promotes sensitivity to irinotecan and rabeprazole prevents this effect, resulting in drug resistance. To ensure the best chance at effective treatment, rabeprazole may not be a suitable PPI for cancer patients treated with irinotecan.

## Introduction

Topoisomerase I (topoI) was identified as a specific target for camptothecin (CPT) and its analogues like irinotecan[1]. Irinotecan is frequently used to treat colon, ovarian, pancreatic, breast, and small cell lung cancer. TopoI reduces DNA supercoiling by cutting and re-ligating DNA, and a controlled rotation between the cutting and re-ligation cycles reduces the supercoiling. However, in the presence of irinotecan, the re-ligation cycle is inhibited, and collision with a replication fork leads to a DNA double strand break (DNA-DSB)[2]. The topoI inhibitor camptothecin and its analogues (CPTs), like irinotecan and topotecan are used extensively to treat various solid tumors. However, only 13-32% of patients respond, and the mechanisms of resistance are not well-understood[3]. Classical mechanisms of irinotecan resistance potentially involve an ATP-binding cassette (ABC) transporter, ABCG2, which reduces intracellular drug accumulation [4–6]. Inhibition of the ABCG2 drug efflux pump using sorafenib sensitizes both non-resistant cells and resistant cells to irinotecan[7, 8]. Another proposed mechanism for irinotecan resistance is topoI gene mutation[9]. However, DNA sequencing studies have failed to find topoI gene mutations in cancer patients[10, 11]. Other mechanisms may include faster repair of trapped topoI protein-linked DNA breaks, which is associated with perturbed histone acetylation[8]. However, none of these studies have been conclusively shown to impart cellular resistance to irinotecan. In response to irinotecan, topoI is ubiquitinated and degraded by ubiquitin proteasomal pathway (UPP) and rapid topoI degradation was shown to be associated with irinotecan resistance[12]. Recently we have published the mechanism of topoI degradation by UPP, and have shown that the DNA-PKcs kinase cascade determines the rate of topoI degradation. Importantly, deregulated kinase cascade keeps the DNA-PKcs constantly active, leading to higher basal levels of phosphorylated topoI-S10, rapid degradation of topoI, and irinotecan resistance. To identify the upstream regulator of DNA-PKcs, we performed a siRNA library screen of 56 reported nuclear phosphatases, and subsequently phosphatase-silenced cells were treated with irinotecan to determine the CPT induced rate of topoI degradation. Carboxy-terminal domain RNA polymerase II polypeptide small phosphatase 1 (CTDSP1) was identified as one of the strongest upstream regulator of DNA-PKcs dependent proteasomal degradation of topoI. The DNA-dependent protein kinase (DNA-PK) is a serine/threonine protein kinase composed of a large catalytic subunit (DNA-PKcs) and Ku 70/80 heterodimer[13]. It is now established that the Ku 70/80 heterodimer binds broken DNA double strands and recruits DNA-PKcs in vitro[14]. Once recruited, DNA-PK stabilizes the broken strand, targets Artemis for end processing, and finally the ligation is carried out by the XLF/XRCC4/Lig IV complex. This classic NHEJ is the major eukaryotic pathway for DNA double strand break repair[15]. Although DNA-PKcs is considered a component of DNA damage response, recent findings indicate a variety of other important roles in genome maintenance and tumor pathogenesis[16]. However, our understanding of the role of DNA-PKcs in tumor pathogenesis is not understood, and regulation of kinase activity is partly understood. Autophosphorylation at multiple S/T sites is the first mechanism of DNA-PKcs kinase activation and inactivation particularly in response to DNA-DSB. The second regulatory component is dephosphorylation of DNA-PKcs by phosphatases[13].

DNA-dependent RNA polymerase II consists of 12 polypeptides. The largest polypeptide Rpb1 contains heptad repeats, (Tyr-Ser-Pro-Thr-Ser-Pro-Ser) in the c-terminus domain (CTD), this unique feature of RNAPII distinguishes it from other polymerases[17]. The 52-heptad tandem repeats of CTD determine the regulation of transcription. Post-translational modifications of the 7 amino acid tandem repeats, particularly Ser2 and Ser5, are considered critical in RNAP II - dependent mRNA transcription. The pattern of CTD phosphorylation during the transcription cycle is highly dynamic and requires the activity of dedicated phosphatases and kinases[17]. Several kinases have been identified to phosphorylate CTD, most notably CDK7, CDK8, and CDK9. FCP1, Ssu72, and small CTD phosphatases (SCP) dephosphorylate the CTD during the different phases of MRNA transcription. CTDSP1 is part of the SCP (Small CTD Phosphatases) family, whose loss of function results in tumor development and proliferation[18]. CTDSP1 is a nuclear phosphatase that associates with RNAP II and de-phosphorylates the serine 5 of the heptad repeat[19, 20]. The phosphorylation state of RNAP II affects its transcription activity; therefore CTDSP1 plays a role in gene expression regulation[20]. Recent studies have shown that the activity of CTDSP1 is suppressed by the proton pump inhibitor, rabeprazole[21].

In this study, we demonstrated that CTDSP1 promotes sensitivity to irinotecan and rabeprazole prevents this effect, resulting in drug resistance.

## Materials and Methods

### Cell culture and drug treatment

The human colon cancer cell lines HCT116, HT29, DLD1, and LoVo were obtained from ATCC. HCT116 and DLD1 cells were grown and maintained in RPMI containing 10% FBS and pen-strep. HT29 cells were grown and maintained in McCoy’s medium containing 10% FBS and pen-strep. LoVo cells were grown and maintained in Ham’s F12-K containing 10% FBS and pen-strep. All cells were grown at 37 °C and 5% CO_2_ in a humidified cell culture incubator. Topo I inhibitor treatment was performed using various concentrations of either irinotecan (Sigma-Aldrich) or SN-38 (Tocris). SN-38 is the irinotecan active metabolite. Cells were also treated with various concentrations of Rabeprazole (Santa Cruz Biotechnology).

### CTDSP1 siRNA transfection

CTDSP1-specific siRNA (Silencer Predesigned siRNA: 5’-CCUCGUGGUUUGACAACAU-3’) and negative control were purchased from Dharmacon. Transfection of HCT116 cells (0.6×10^6^ cells/well in 6-well plate) with siRNA oligonucleotides was performed using Lipofectamine RNAiMAX (Invitrogen).

### CTDSP1 overexpression

For reverse transcription quantitative RT-PCR, the following mRNA sequences were used: CTDSP1, forward primer, 5’-CACCATGGACAGCTCGGCCGTCATTACTC-3’, and reverse primer, 5’-CTAGCTCCCTGGCCGTGGCTGCCTG-3’. The cDNA was synthesized from total RNA of HT29 cells using Super Script 3 First-Strand Synthesis (Invitrogen). Quantitative PCR amplification was performed using a C1000 Touch Thermal Cycler. The PCR product and the pENTR/D-TOPO vector were mixed and incubated for 5 min at room temperature. The reaction was used to transform competent *E. coli* cells (Mach1, Invitrogen). The recombinant bacteria were screened using LB agar medium containing kanamycin. The recombinant plasmid pENTR-CTDSP1 was extracted from positive colonies using the QIAprep Spin Miniprep Kit (Qiagen) and proper orientation of the cloned fragment was verified by PCR and DNA sequencing. The recombinant plasmid was transfected into HT29 cells using 4D-Nucleofecor (Lonza).

### Immunoblotting

Cells cultured in 6-well plates were scraped into an ice-cold RIPA buffer. Samples were clarified by centrifugation at 15000 rpm for 15 min at 4°C. We used the iBind Western System (Invitrogen) and performed the imaging using the Amersham Imager 600 instrument (GE Healthcare, Little Chalfont, UK). Primary antibodies included antibodies against β-actin (Cell Signaling Technology), CTDSP1 (Abcam), Topoisomerase I (BD Pharmingen), and p-DNAPKcs (Abcam). Antigen/antibody complexes were visualized by enhanced chemiluminescence (ECL detection system).

### Immunofluorescence

Control and CTDSP1 knockdown cells were grown on sterile coverslips in 100mm plates and after washing with PBS, cells were fixed with 3.7% formaldehyde. After 25 minutes of fixation, coverslips were washed with 0.2% Triton-X-100 and then blocked with 3% BSA for 1 hour. After blocking, each coverslip was incubated with a monoclonal anti-phosphorylated DNAPK antibody, followed by Alexa-fluor 488-conjugated goat anti-rabbit IgG. Nuclei were stained with DAPI and imaging was performed on the Leica SP5 fluorescence microscope.

### Integrating EGFP following the *hTOPI* gene in HCT-116 cells

Approximately 1000 bases of the genomic sequence flanking the *hTOPI* exon in HCT116 cells were sequenced to identify cell specific polymorphisms. PCR products across the genomic sequence were amplified and then cloned with the Zero Blunt TOPO PCR Cloning Kit (Invitrogen), and then transformed into E. coli XL-1 blue competent cells. Approximately 25 colonies were grown overnight in a 37°C incubator prior to Sanger sequencing and miniprep. Designed single guide-RNA (sgRNA) targeting *hTOPI* stop codon, such that the SpCas9 binding site would be eliminated after gene conversion. Areas of the *hTOPI* stop codon and genomic regions flanking it were assembled with the EGFP-P2A-Pac fusion cassette for puromycin resistance to create the final donor plasmid. HCT-116 cells were transfected using the lipofectamine method (Life-technology). This method utilizes: 2μg of wt *Streptococcus pyogenes* Cas9 plasmid, 1μg of the sgRNA plasmid, and 2μg of the homologous recombination donor plasmid. Cells were selected for with 4μg/mL puromycin after 7 days of transfection. Cells were sorted by the MoFlo Legacy (Beckman Counter) and grown for 14 days for selection of genomically edited single cell clones. Cells were then maintained in puromycin -treated media. Incorporation was confirmed using PCR amplification of the genomic locus.

### Cell viability assay

Cells were seeded in 96-well plates at 700-2000 per well in 180 μl of medium. Next, a 100 μM SN-38 stock solution was used to prepare dilutions in 20 μl of medium before treatment. After incubating for 72 h, cells were incubated CellTiter-Glo 2.0 reagent into each well. Cell viability was then measured by luminescence detection using a Synergy H1 microplate reader (BioTek).

### Clonogenic assay

All cells were treated with various concentrations of SN-38 (0, 2.5, 5.0, or 7.5 nM) for 24 h, and then 50 cells per well were seeded in 6-well plates. After 14 days, when colonies were apparent, cells were fixed in 6% glutaraldehyde and 0.5% crystal violet for 15 min, then washed with tap water. The colonies per well were counted.

### Patients

Retrospective data from 61 patients who underwent 2^nd^-line chemotherapy including irinotecan between January 2003 and December 2015 at the Department of Surgery and Science, Kyushu University, were retrieved. Of the 61 patients, 5 patients who discontinued chemotherapy or whose data was not complete, were excluded from the study. Finally, 56 patients were eligible for analysis.

### Study approval

All experiments were conducted with approval by the Ethics Committee of the Kyushu University (Approval Number: No. 30-215)

## Results

### CTDSP1 expression correlated with topo I degradation in colorectal cancer cell line

We have previously reported that topo I is phosphorylated by DNA-PKcs at serine-10, and the resulting phosphoprotein (topoI-pS10) is efficiently ubiquitinated and targeted for rapid proteasomal degradation[3]. Protein phosphatases have been shown to interact with DNA-PKcs and could serve as upstream regulators of DNA-PKcs. To identify potential upstream regulators of DNA-PKcs, we screened a siRNA library comprising all human nuclear phosphatases. Our data demonstrated that silencing of CTDSP1 significantly altered the rate of topoI degradation. In the present study, the possible role of CTDSP1 as a regulator of topoI stability and irinotecan sensitivity was tested first by assessing the relationship between CTDSP1 expression and rate of topoI degradation. Cell lysates from four CRC cell lines, HCT116, DLD1, LoVo, and HT29 were subjected to immunoblot analysis with anti-CTDSP1. The results indicate a higher level of CTDSP1 in HCT116 cells and the lowest level in HT29 (Fig 1A). HCT116 and HT29 cells were treated with SN-38 for 90 and 180 min and cell lysates were subjected to determine the rate of topoI degradation. The lower level of topoI protein at 180 min clearly indicates a higher rate of topoI degradation in HT29 cells compared to HCT116 cells (Fig 1B). We then asked the obvious question: does this SN-38-induced rate of topo I degradation have an impact on drug sensitivity? The clonogenic survival assay clearly demonstrated that HT29, with low CTDSP1 and rapid topo I degradation, is more resistant to HCT116 cells (Fig 1C).

**Fig 1.**
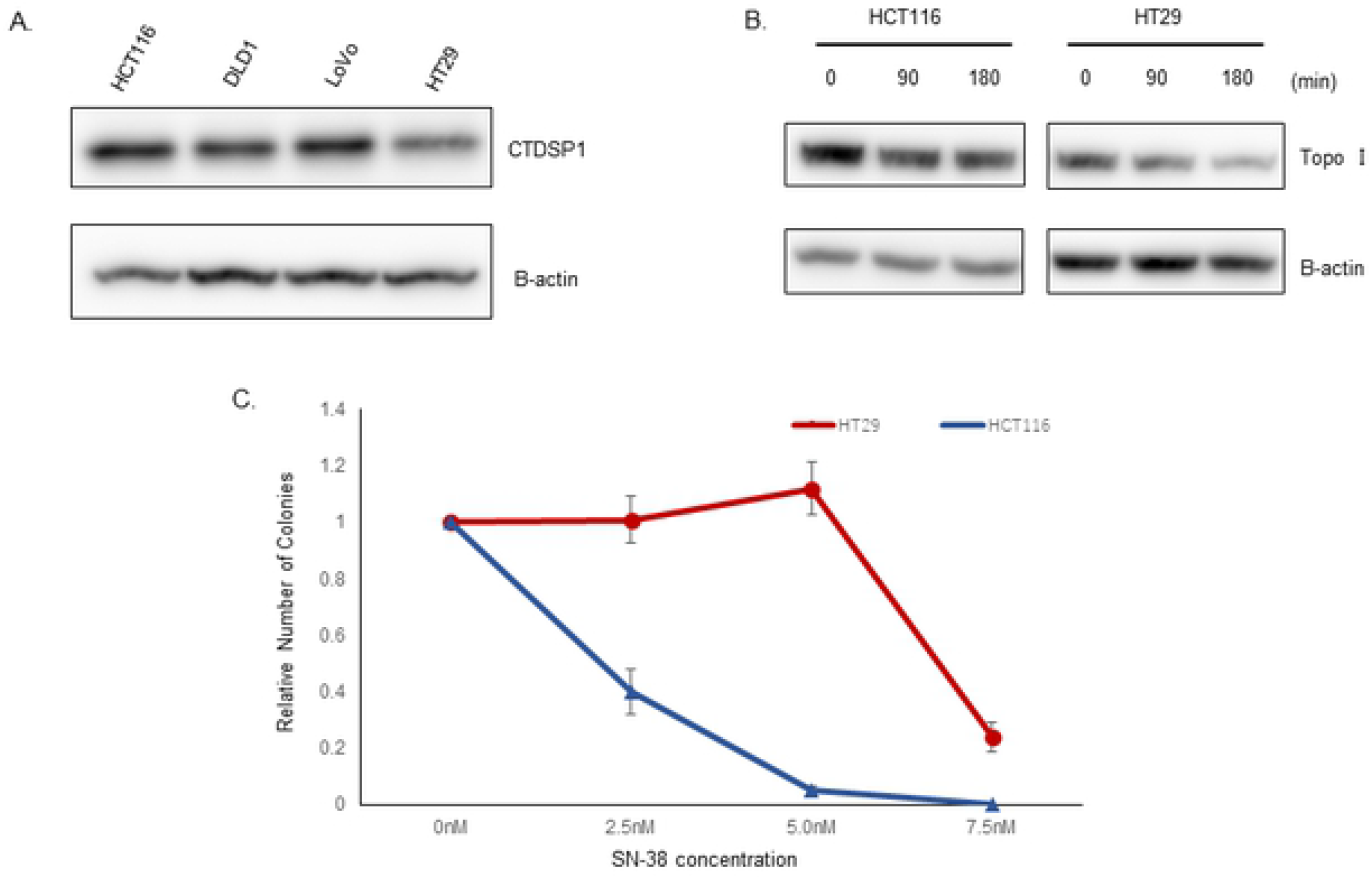
CTDSP1 regulates irinotecan sensitivity in colon cancer cell lines. **A**, Cell lysates from four colon cancer lines, HCT15, DLD1, LoVo and HT29, were immunoblotted with anti-CTDSP1 and anti-β-actin antibodies. **B**, HCT116 and HT29 cells were treated with 2.5 μM SN-38 and harvested after 90 and 180 min. Cell lysates were immunoblotted with anti-topo I and anti-β-actin antibodies. **C**, HCT116 and HT29 cells were treated with various concentrations of SN-38 and clonogenic assays were performed to determine the relative number of colonies.

### Silencing of CTDSP1 enhances topoI degradation and irinotecan resistance

Higher levels of CTDSP1 negatively regulate DNA-PKcs-dependent topoI phosphorylation and rate of topoI degradation. We silenced CTDSP1 by siRNA and asked if topo I degradation can be enhanced and cellular resistance to SN-38 can be induced. Post silencing, a significant reduction in CTDSP1 protein level was observed in HCT116 cells compared to HCT116 control cells (Fig 2A). The CTDSP1-deficient cells were treated with SN-38 for 90 and 180 minutes. Immunoblot analysis of the cell lysates clearly demonstrated a higher rate of topoI degradation in cells that were deficient in CTDSP1 (Fig 2B). This was observed both by immunoblot analysis (Fig 2C). The si CTDSP1 cells were also treated with various concentrations of SN-38, and cell survival was compared with control siHCT116 cells. The data clearly demonstrated that siCTDSP1-HCT116 cells are significantly more resistant to SN-38 (Fig 2D). Similarly, a clonogenic assay demonstrated that siCTDSP1-HCT116 are resistant to SN-38 (Fig 2E).

**Fig 2.**
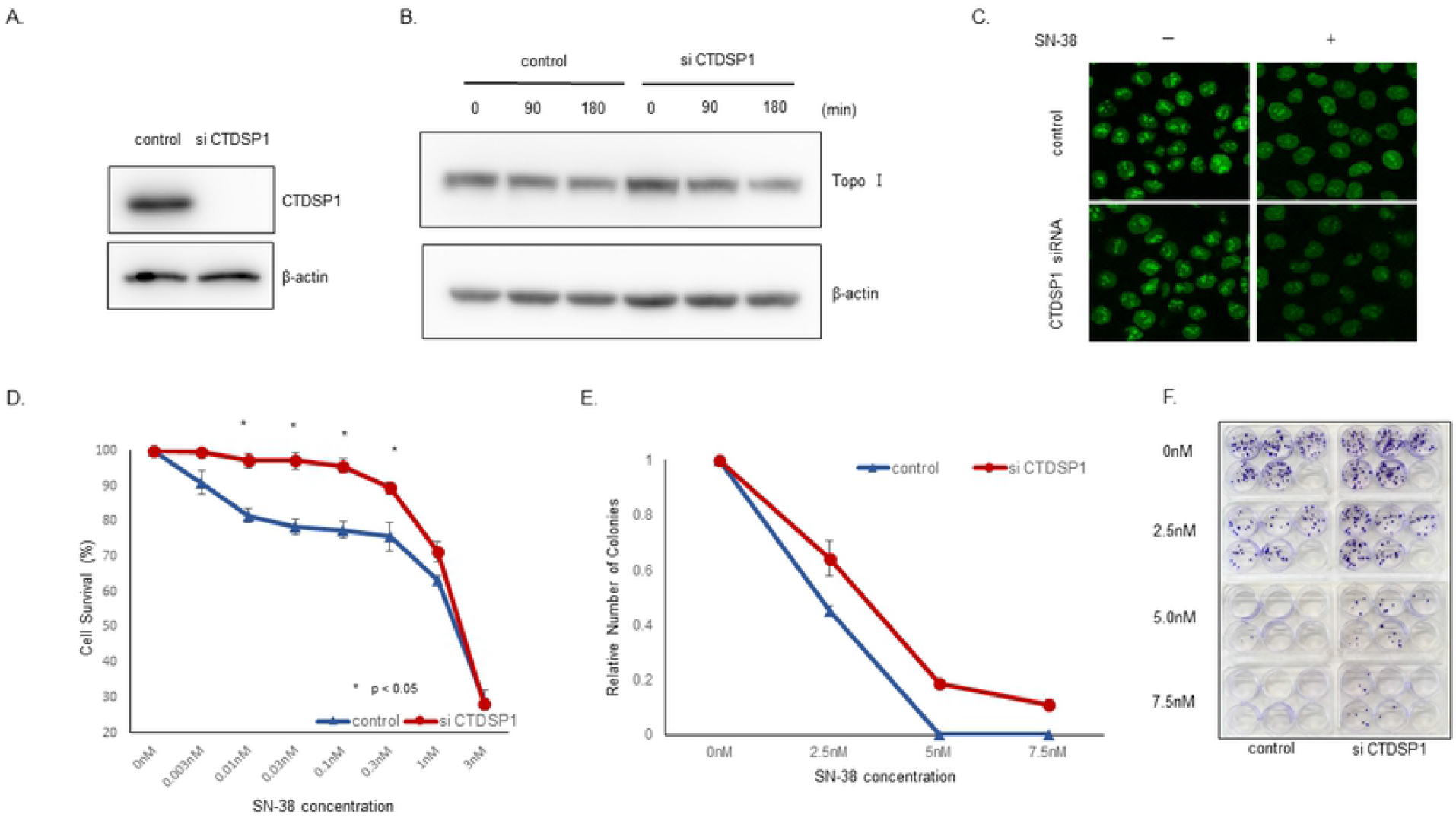
Silencing of CTDSP1 enhances topoI degradation and irinotecan resistance. **A**, Cells transfected with CTDSP1 or control siRNA were lyzed and the cells lysates were immunoblotted with anti-CTDSP1 and anti-β-actin antibodies. **B**, HCT116-siRNA CTDSP1 or control siRNA, treated with 2.5 μM SN-38 were harvested after 90 and 180 min. Cell lysates were immunoblotted with anti-topo I and anti-β-actin antibodies. Cells lysates were immunoblotted with anti-topo I and anti-β-actin antibodies. **C**, EGFP was integrated with topo I in HCT116 cells using CRISPR/Cas9 system and CTDSP1 was knocked down in this cell line by siRNA. Cells were treated with 2.5uM SN-38 for 60 min and the topo I-EGFP signal was imaged by Leica SP5 confocal microscope. **D**, HCT116-siRNA-CTDSP1 or control siRNA were plated in a 96-well plate and treated with various concentrations of SN-38 for 72 h. Cell viability was determined by detecting the luminescence. **E**, HCT116-siRNA-CTDSP1 or control siRNA cells were plated in a 6-well plate and treated with various concentrations of SN-38 for 24 h. Then, 50 cells per well were plated in a 6-well plate. After 14 days, when colonies were apparent (right panel), colonies were counted and relative number of colonies was determined (left panel).

### The higher expression of CTDSP1 in HT29 cells inhibits topoI degradation and restores SN-38 sensitivity

CTDSP1 protein level was significantly low in HT29 cells. We increased the expression of CTDSP1 in these cells and asked if the SN-38-induced topoI degradation would be altered. HT29 cells were transfected and selected for CTDSP1 expression. A significantly higher level was observed in overexpressing cells (Fig 3A). The Vector control and CTDSP1-expressing HT29 cells were then treated with SN38, and topoI protein level was analyzed after 90 and 180 min. The data clearly demonstrates that in cells with higher CTDSP1, the topoI degradation was minimal. In contrast, a significantly lower level of topoI was observed in HT29-vector cells. This indicates that overexpression of CTDSP1 in HT29 regulates SN-38-induced proteasomal degradation of topoI (Fig 3B). The cell viability assay indicated that topoI stabilization by CTDSP1 overexpression restores SN-38 drug sensitivity (Fig 3C).

**Fig 3.**
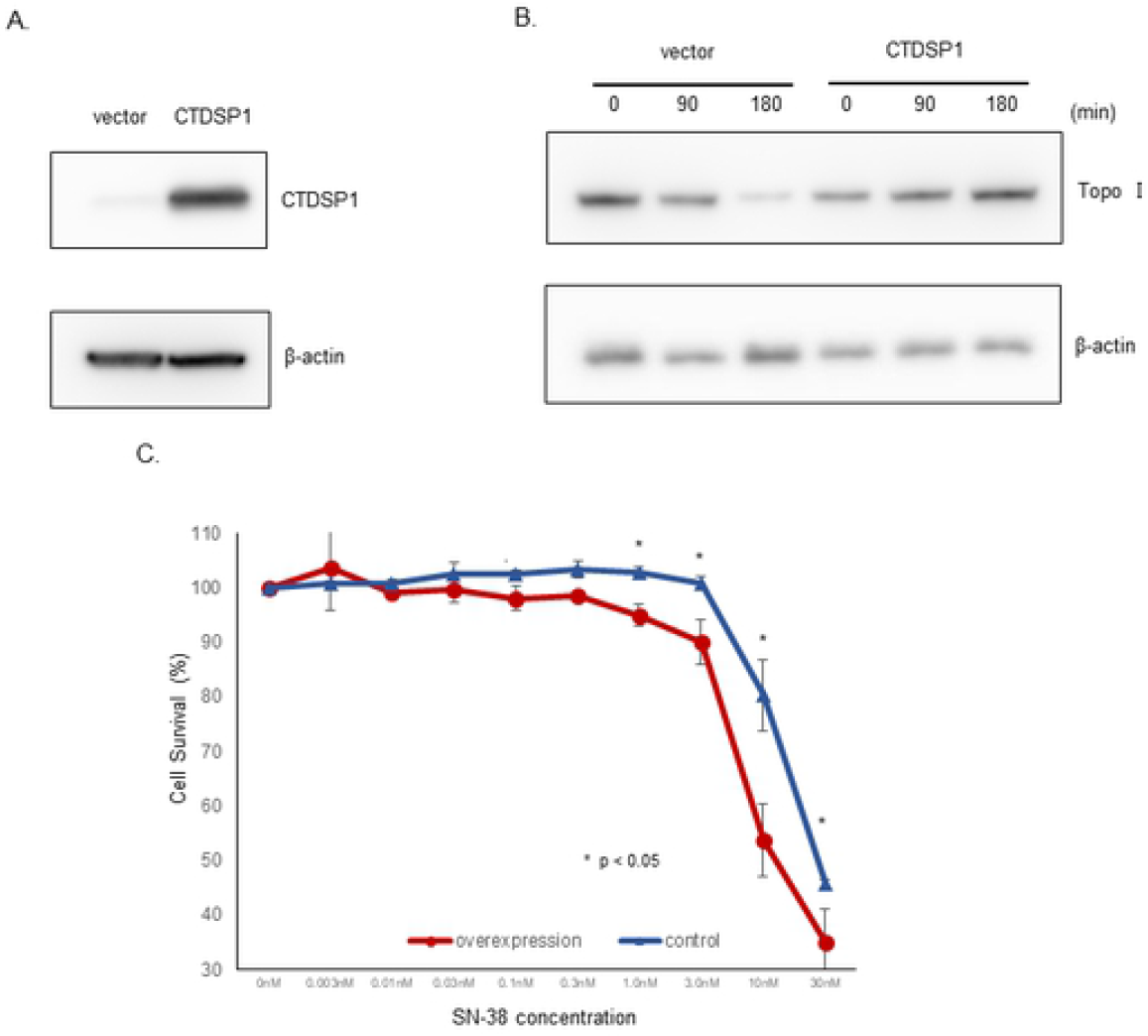
CTDSP1 restores irinotecan sensitivity in HCT116 cells. **A**, CTDSP1-overexpressing and control cells were lyzed and the cells lysates were immunoblotted with anti-CTDSP1 and anti-β-actin antibodies. **B**, CTDSP1-overexpressing and control cells were treated with 2.5 μM SN-38 and harvested after 90 or 180 min. Cells lysates were immunoblotted with anti-topo I and anti-β-actin antibodies. **C**, CTDSP1-overexpressing and control cells were plated in a 96-well plate and treated with various concentrations of SN-38 for 72 h. Cell viability was determined by luminescence detection.

### CTDSP1 activates DNA-PKcs and enhances topo I degradation

Once we determined that HCT116 and HT29 have differential expression of CTDSP1, we asked if DNA-PKcs activation status is different in these two-cell lines. Cell lysates from these cells were subjected to immunoblot analysis with anti-DNA-PKcs-pS2056. The results indicated a higher level of phosphorylated DNA-PKcs in HT29 cells (Fig 4A). In contrast, a minimal phosphorylation of DNA-PKcs was observed in HCT116 cells. We asked if silencing of CTDSP1 would have an impact on the activation status of DNA-PKcs. Cell lysates from si-CTDSP1-HCT116 cells were analyzed by immunoblotting, and the data clearly demonstrated that silencing of CTDSP1 significantly enhanced phosphorylated DNA-PKcs in HCT116 cells (Fig 4B). This was observed by GFP florescence level by confocal microscopy (Fig 4C).

**Fig 4.**
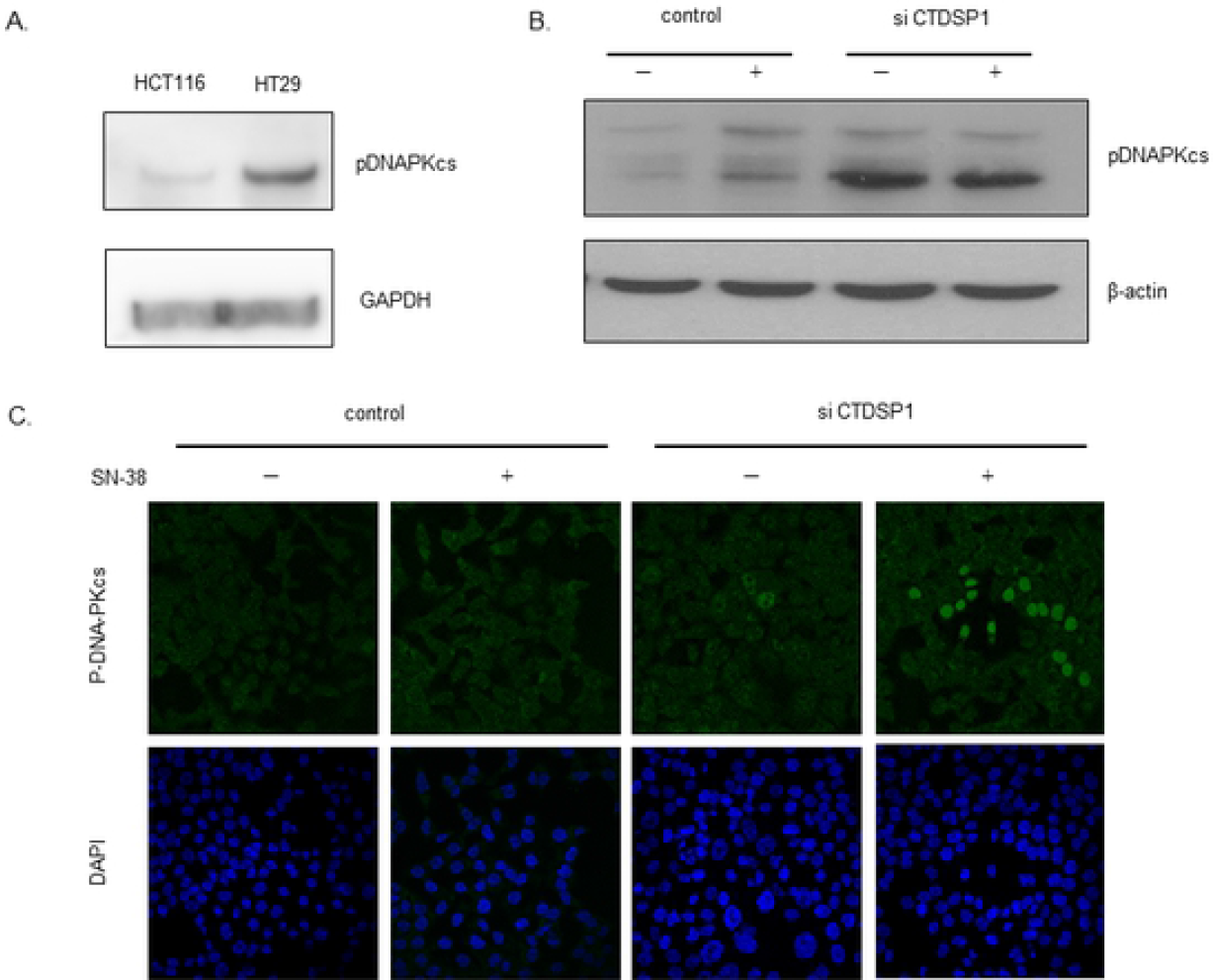
CTDSP1 activates DNA-PKcs and enhanced DNA-PKcs dependent topoI degradation in response to irinotecan. **A**, HCT116 and HT29 cell lysates were immunoblotted with anti-pDNAPKcs and anti-GAPDH antibodies. **B**, CTDSP1 was silenced by CTDSP1 siRNA in HCT116 cells and control and CTDSP1 silenced cell lysates were subjected to immunoblot with anti-CTDSP1 and anti-β-actin. **C**, Control and CTDSP1 knockdown cells were treated with SN38 and analyzed by immunofluorescence staining with anti-p-DNA-PKcs (S2056). The cells were analyzed by Leica SP5 confocal microscope.

### Rabeprazole inhibits CTDSP1 and enhances topoI degradation and irinotecan resistance

Rabeprazole binds to the hydrophobic pocket of CTDSP1 and inhibits its phosphatase activity. This hydrophobic pocket is adjacent to the active site in CTDSP1 and rabeprazole most likely acts as a direct competitor of the natural substrate, i.e., the CTD phosphorylated peptide[21]. We treated HCT116 cells with rabeprazole and then with SN-38. Cells were harvested at 90, 180, and 270 min post-SN-38 treatment, and cell lysates were analyzed to determine the rate of topo I degradation. The results demonstrated that topoI was degraded at 270 min (Fig 5A). To visualize the rate of topoI degradation, HCT116-topoI-GFP cells were treated first with rabeprazole, and then treated with 5 and 10 μM SN-38 for 60 min (Fig 5B). A significant reduction of topoI-GFP florescence at 10 μM rabeprazole at 60 min clearly indicated a rapid rate of topoI degradation in cells that were treated with rabeprazole. To understand if rabeprazole-mediated inhibition of CTDSP1 negatively regulates DNA-PKcs, rabeprazole-treated cells were treated with SN-38 and cell lysates were analyzed at 90 and 180 min. Anti-DNA-PKcs-pS2056 immunoblot analysis clearly demonstrated significant kinase activation at 90 min post-SN-38 treatment (Fig 5C). This was observed by GFP florescence level by confocal microscopy (Fig 5D). This data clearly demonstrated that rabeprazole, a specific CTDSP1 inhibitor, activates DNA-PKcs and enhances SN-38-induced topoI degradation in HCT116 cells. We asked if rabeprazole-mediated inhibition of CTDSP1 also induces SN-38 resistance in HCT116 cells. Cell survival and clonogenic assays were performed and significant increases in drug resistance were observed in both assays (Fig 5E, F)

**Fig 5.**
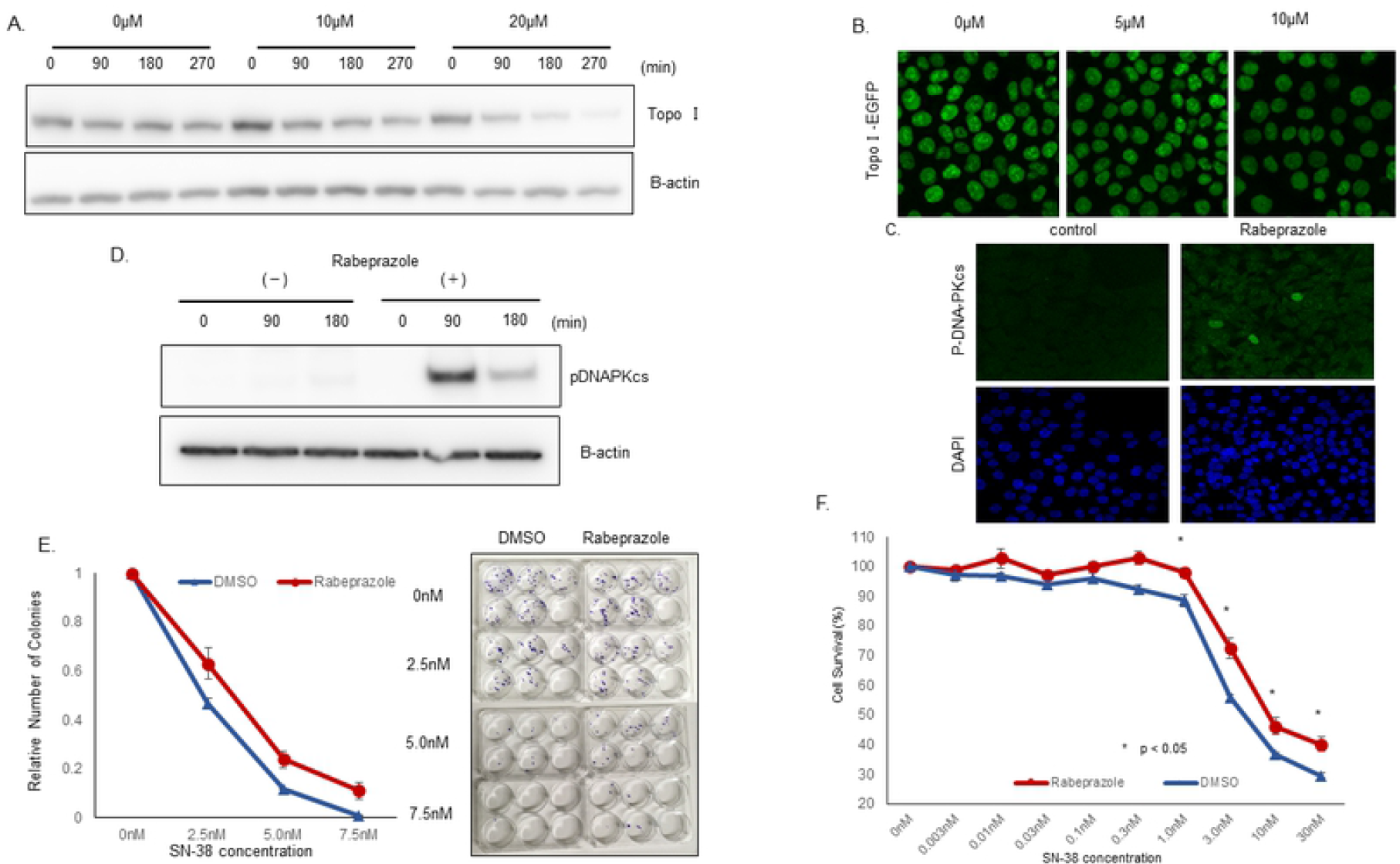
Rabeprazole promotes topo I degradation and irinotecan resistance. **A**, HCT116 cells were plated in a 6-well plate and treated with various concentrations of rabeprazole (0, 10, 20 μM) for 72 h, and then with 2.5 μM SN-38 and harvested after 90 or 180 min. Cells lysates were immunoblotted with anti-topo I and anti-β-actin antibodies. **B**, Genomically edited HCT116 cells with TopoI-EGFP fusion protein were treated with 5 and 10 μM of Rabeprazole for 48 hours and topoI-GFP protein level was analyzed by confocal microscope. **C**. HCT116 cells were plated in a 6-well plate, treated with 40 μM rabeprazole or DMSO for 72 h, and then with 2.5 μM SN-38, and harvested after 90 or 180 min. Cell lysates were immunoblotted with anti-pDNA-PKcs and anti-β-actin antibodies. **D**, HCT 116 cells were treated with rabeprazole, control and treated cells were analyzed by immunofluorescence analysis with anti-phospho-DNA-PKcs (S2056) and confocal microscopy. **E**. HCT116 cells were plated in a 6-well plate and treated with rabeprazole or DMSO for 72 h. Then, 50 cells were plated in each well of 6-well plate and treated with various concentrations of SN-38 for 24 hours. Cell colonies were counted after 14 days. **F**. HCT116 cells were plated in a 6-well plate and treated with 40 μM rabeprazole or DMSO for 72 h. Then, cells were plated in a 96-well plate and treated with various concentrations of SN-38 for 72 h. Cell viability was determined by luminescence detection.

### Treatment with rabeprazole may reduce the effect of irinotecan in colorectal cancer patients

We next examined clinical data to investigate whether rabeprazole induced irinotecan resistance in patients with colon cancer. Of 56 eligible patients who underwent chemotherapy with irinotecan, 50 (89.3%) had not received rabeprazole during or before chemotherapy, while 6 (10.7%) had been treated with rabeprazole (Fig 6). We compared the overall response in the two groups. Partial response (PR) was observed in only one patient (16.7%) of the rabeprazole group (Table 1). CR and PR were observed in 1 (2.0%) and 14 patients (28.0%), respectively, of the non-rabeprazole group. This result suggested that rabeprazole had an effect on the response to irinotecan in clinical settings.

**Fig 6.**
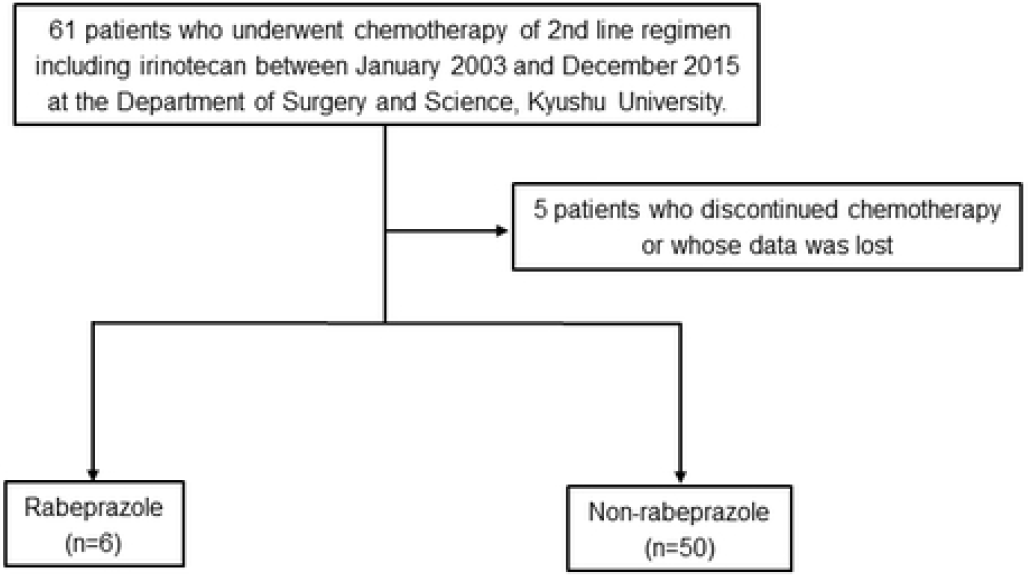
Flow chart depicting the process of patient selection.

**Table 1.**
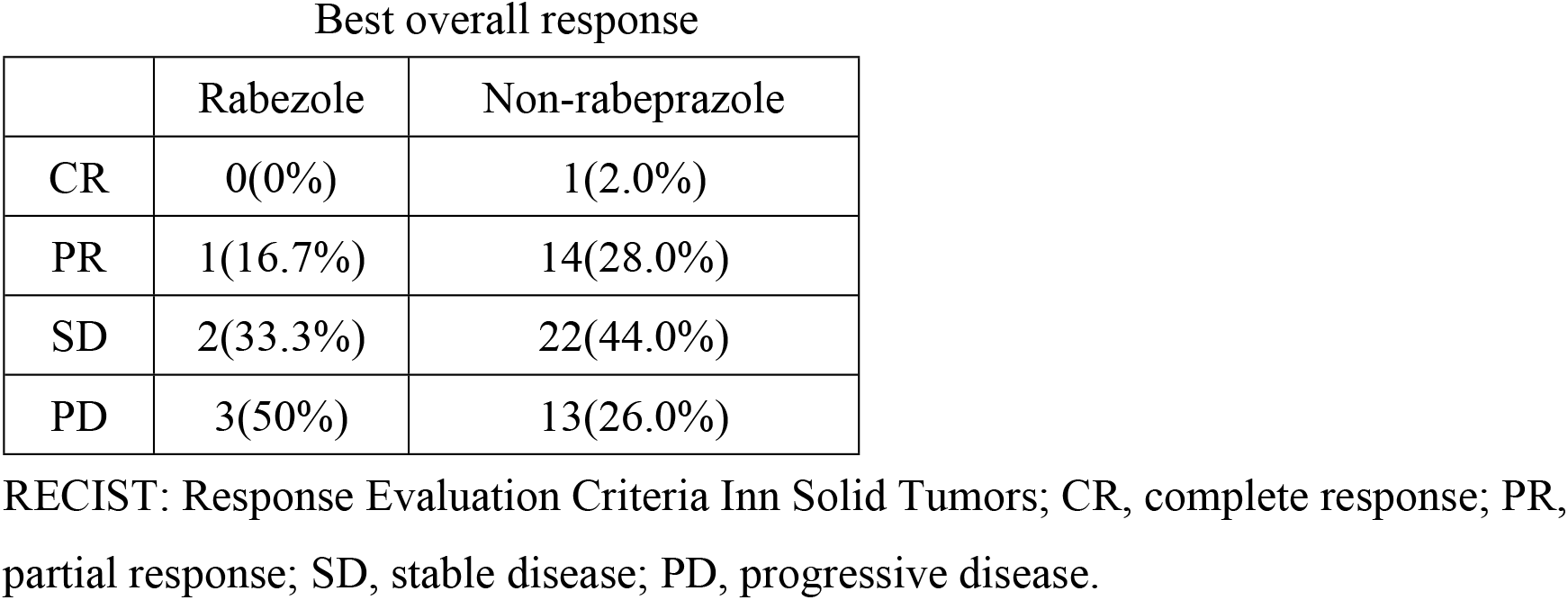
The overall response in the rabeprazole and non-rabeprazole group

| Best overall response |  |  |
| --- | --- | --- |
|  | Rabazole | Non-rabeprazole |
| CR | 0(0%) | 1(2.0%) |
| PR | 1(16.7%) | 14(28.0%) |
| SD | 2(33.3%) | 22(44.0%) |
| PD | 3(50%) | 13(26.0%) |

RECIST: Response Evaluation Criteria Inn Solid Tumors; CR, complete response; PR, partial response; SD, stable disease; PD, progressive disease.

## Discussion

One of the most remarkable phenomena observed in the cellular response to irinotecan is degradation of topoI. It was first observed in leucocytes of patients treated with 9-nitro-camptothecin in phase 1 clinical trials. The leucocytes isolated after 24, 48, and 72 hours demonstrated decreasing topoI protein levels[22]. Later, these findings were reproduced in cancer cells[23]. More importantly, Desai et al, in a series of publications, demonstrated that the irinotecan-induced degradation of topoI is mediated by a ubiquitin proteasomal pathway, cells degrade topoI differentially, and the cells that degrade topoI rapidly are resistant to irinotecan[24]. More recently we have demonstrated that: i) topoI associates with DNA-PK, and DNA-PKcs phosphorylates topoI at Serine 10; ii) Phosphorylated topoI is ubiquitinated by BRCA1; iii) cells with higher basal levels of topoI-pS10 degrade topoI rapidly and are resistant to irinotecan; iv) the higher basal level of topoI-pS10 is maintained by phosphatase dependent activation of DNA-PKcs v) nuclear phosphatase siRNA library screen identified PTEN and CTDSP1 enhances irinotecan induced topoI degradation and vi) silencing of PTEN enhanced DNA-PKcs activity and irinotecan resistance[3]. Phosphorylation dependent activation and inactivation of DNA-PKcs is well documented. Autophosphorylation of several S/T moieties localized in ABCD and PQR clusters is at the core of regulation of DNA-PKcs kinase activity in response to DNA-DSB and phosphorylation of serine 2506 indicates the activation of DNA-PKcs[13]. The second proposed mechanism of DNA-PKcs regulation depends on the dephosphorylation by phosphatases. In one of the early reports, protein phosphatase 1 or protein phosphatase 2A were shown to reactivate autophosphorylated inactive DNA-PKcs. Furthermore, the addition of a phosphatase inhibitor reverses this process[25]. PP2A was also shown to play a role in NHEJ by directly dephosphorylating DNA-PKcs[26]. Another phosphatase, PP6, was shown to regulate the kinase activity of DNA-PKcs in response to DNA-DSB. Both catalytic units (PP6C) and regulatory subunits (PPR1) of PP6 interact with DNA-PKcs, and silencing of PP6C induced IR sensitivity and delayed release from the G2/M checkpoint[27, 28].

Our siRNA library screen of nuclear phosphatase identified PTEN and CTDSP1 as two phosphatases that significantly enhance the CPT induced topoI degradation. Importantly, we have demonstrated that PTEN regulates DNA-PKcs kinase activity in this pathway and PTEN deletion ensures DNA-PKcs dependent higher topoI-pS10, rapid topoI degradation and irinotecan resistance[3]. We asked if CTDSP1 also dephosphorylates DNA-PKcs, and is the upstream regulator of CPT induced topoI degradation. CTDSP1 is a SCP1 family of phosphatases that dephosphorylate RNAP II and plays a regulatory role in mRNA transcription. Our novel finding indicates that CTDSP1 dephosphorylates DNA-PKcs, changes its kinase activity and regulates irinotecan induced topoI degradation. Our library screen demonstrated that silencing of CTDSP1 enhances irinotecan induced topoI degradation in HCT15 cells. HCT15 cells degrade topoI rapidly and are resistance to irinotecan. To determine the activation of this pathway in cells that do not degrade topoI we used WT and genomically edited HCT116 cells. We impaired CTDSP1 phosphatase activity either by silencing by siRNA or using a specific inhibitor. The results clearly demonstrated that either CTDSP1 inhibition or silencing caused the activation of DNA-PKcs, indicated by phosphorylation of DNA-PKcs-S-2056. Importantly, the knocking down of CTDSP1 resulted in higher basal phosphorylation of DNA-PKcs-S-2056. Similar results were observed when cells were treated with rabeprazole. The activated state changed in response to irinotecan when DNA-PKcs-S2056 protein level was analyzed by immunoblotting and immunofluorescence. One of the important physiological functions attributed to DNA-PKcs in irinotecan-induced pathways is the degradation of topoI. HCT116 cells do not degrade topoI, however when CTDSP1 function was inhibited a rapid topoI degradation was observed. Immunoblot analysis of topoI and reduction in the florescence intensity of topoI-EGFP clearly demonstrated enhanced topoI degradation in CTDSP1 deficient cells. Others and we have shown that enhanced rates of topoI degradation cause irinotecan resistance. To further validate this pathway, we asked if CTDSP1 mediated activation of DNA-PKcs and rapid degradation of topoI causes irinotecan resistance. Cells that were treated with rabeprazole did show irinotecan resistance in both cell survival and clonogenic assay.

CTDSP1 is a member of phosphatases that rely on the DxDx motif and Mg^2+^ to catalyze the phosphoryl-transfer[20, 29]. Rabeprazole binds to a unique hydrophobic pocket of CTDSP1, located in the insertion domain and adjacent to the DxDx motif. Digestive symptoms are the most feared complications of chemotherapy[30, 31] and a proton pump inhibitor (PPI) like rabeprazole, is frequently prescribed to patients. PPI including rabeprazole was recently reported to enhance the antitumor effects of both docetaxel-cisplatin combinations[32] and 5-FU[33] in cells. PPI can restore drug sensitivity in drug-resistant cells by preventing the increase of extracellular and lysosomal pH and increasing the cytoplasmic retention of doxorubicin[34]. However, another study highlighted a potential adverse effect of PPI on overall survival in colorectal cancer[35]. Treatment with a PPI may result in hypochlorhydria in the stomach, which causes hypersecretion of the hormone gastrin from the gastric antrum[36]. Moreover, *in vitro* studies have shown that hypergastrinemia promotes cell proliferation and migration, and inhibits apotosis[37–39]. The relationship between PPI therapy and chemo-sensitivity is not clear. Notably, none of these studies examined the impact of rabeprazole on irinotecan sensitivity. This study is the first to demonstrate that rabeprazole inhibited CTDSP1 activity and caused resistance to irinotecan. Based on our findings, the concurrent use of rabeprazole should be avoided in cancer patients under treatment with irinotecan. If PPI therapy cannot be discontinued due to the severity of the gastrointestinal symptoms, alternative PPIs should be employed.

This study has some limitations. First, all analyzed patients underwent irinotecan chemotherapy as a second-line regimen. Secondary, this analysis was retrospective. Prospective studies focusing on 1^st^-line irinotecan regimens would be necessary to definitively establish the relationship between rabeprazole and irinotecan.

In conclusion, we showed that CTDSP1 and rabeprazole were involved in irinotecan resistance. This study also indicated that rabeprazole may not be a suitable PPI for cancer patients under treatment with irinotecan.

## Acknowledgments

The authors thank S. Tsurumaru and A. Nakamura for technical assistance with the experiments.

